# Divergent kinase WNG1 is regulated by phosphorylation of an atypical activation sub-domain

**DOI:** 10.1101/2022.03.02.482640

**Authors:** Pravin S. Dewangan, Tsebaot Beraki, Elisia A. Paiz, Delia A. Mensah, Zhe Chen, Michael L. Reese

## Abstract

Apicomplexan parasites like *Toxoplasma gondii* grow and replicate within a specialized organelle called the parasitophorous vacuole. The vacuole is decorated with parasite proteins that integrate into the membrane after trafficking through the parasite secretory system as soluble, chaperoned complexes. A regulator of this process is an atypical protein kinase called WNG1. Phosphorylation by WNG1 appears to serve as a switch for membrane integration. However, like its substrates, WNG1 is secreted from the parasite dense granules, and its activity must therefore be tightly regulated until the correct membrane is encountered. Here we demonstrate that, while another member of the WNG family can adopt multiple multimeric states, WNG1 is monomeric and therefore not regulated by multimerization. Instead, we identify two phosphosites on WNG1 that are required for its kinase activity. Using a combination of *in vitro* biochemistry and structural modeling, we identify basic residues that are also required for WNG1 activity and therefore appear to recognize the activating phosphosites. Among these coordinating residues are the “HRD” Arg, which recognizes activation loop phosphorylation in canonical kinases. WNG1, however, is not phosphorylated on its activation loop, and its activating phosphosites instead appear to lock the kinase C-lobe into an activated conformation. These data suggest a simple model for WNG1 activation by increasing ATP concentration above a critical threshold once WNG1 traffics to the parasitophorous vacuole.

## Introduction

Protein kinases are the largest protein family in eukaryotes and regulate diverse cellular functions by phosphorylation (Hunter, 1995; Manning et al., 2002). To maintain the fidelity of signaling, many kinases must be activated before they can efficiently carry out phosphoryl transfer (Boulton et al., 1991, 1990; Ray and Sturgill, 1988; Robbins et al., 1993; Tao et al., 1970). Indeed, uncontrolled kinase activity is often pathogenic and is therefore at the heart of many diseases (Davies et al., 2002; James et al., 2005; Lacronique et al., 1997; Moelling et al., 1984; Rapp et al., 1983). The two most common mechanisms by which kinase activities are regulated are multimerization (Chao et al., 2010; Oliver et al., 2006; Pike et al., 2008) and phosphorylation on the kinase “activation loop.” (Abe et al., 2001; Canagarajah et al., 1997; Russo et al., 1996; Zheng et al., 1993). While the regulation of canonical protein kinases is now well understood, a subset of protein kinases have a number of atypical motifs and therefore function through unusual mechanisms (Eswaran et al., 2009; Piala et al., 2014; Xu et al., 2002, p. 1; Yang et al., 2015).

Apicomplexan parasites such as *Toxoplasma gondii* encode an unusually high number of pseudokinases and atypical kinases (Peixoto et al., 2010; Talevich et al., 2011), the majority of which are secreted into the host cell during invasion (Bradley et al., 2005; Peixoto et al., 2010; Reese and Boyle, 2012). Among these kinases is a family that lacks the Glycine-rich loop that is required for kinase activity in all other kinases (Beraki et al., 2019; Knighton et al., 1991; Walker et al., 1982). Nevertheless, members of these “<u>W</u>ith-<u>N</u>o-<u>G</u>ly-loop” (WNG) kinases are still able to robustly catalyze phosphoryl transfer using a reorganized kinase structure (Figure 1A). The most highly conserved member of the family, WNG1, is secreted into the *Toxoplasma gondii* parasitophorous vacuole (PV), the organelle in which the parasite grows and replicates within an infected host cell. WNG1 phosphorylates a number of other secreted proteins, and appears to regulate their insertion into the PV membrane (Beraki et al., 2019). Notably, these PV-resident membrane proteins are trafficked through the parasite Golgi and secretory granules as soluble entities (Gendrin et al., 2008), and integrate only into the correct membrane. Because the kinase is secreted from the very same organelles as its substrates, we reason that WNG1 activity must be regulated to ensure correct trafficking of its substrates.

**Figure 1:**
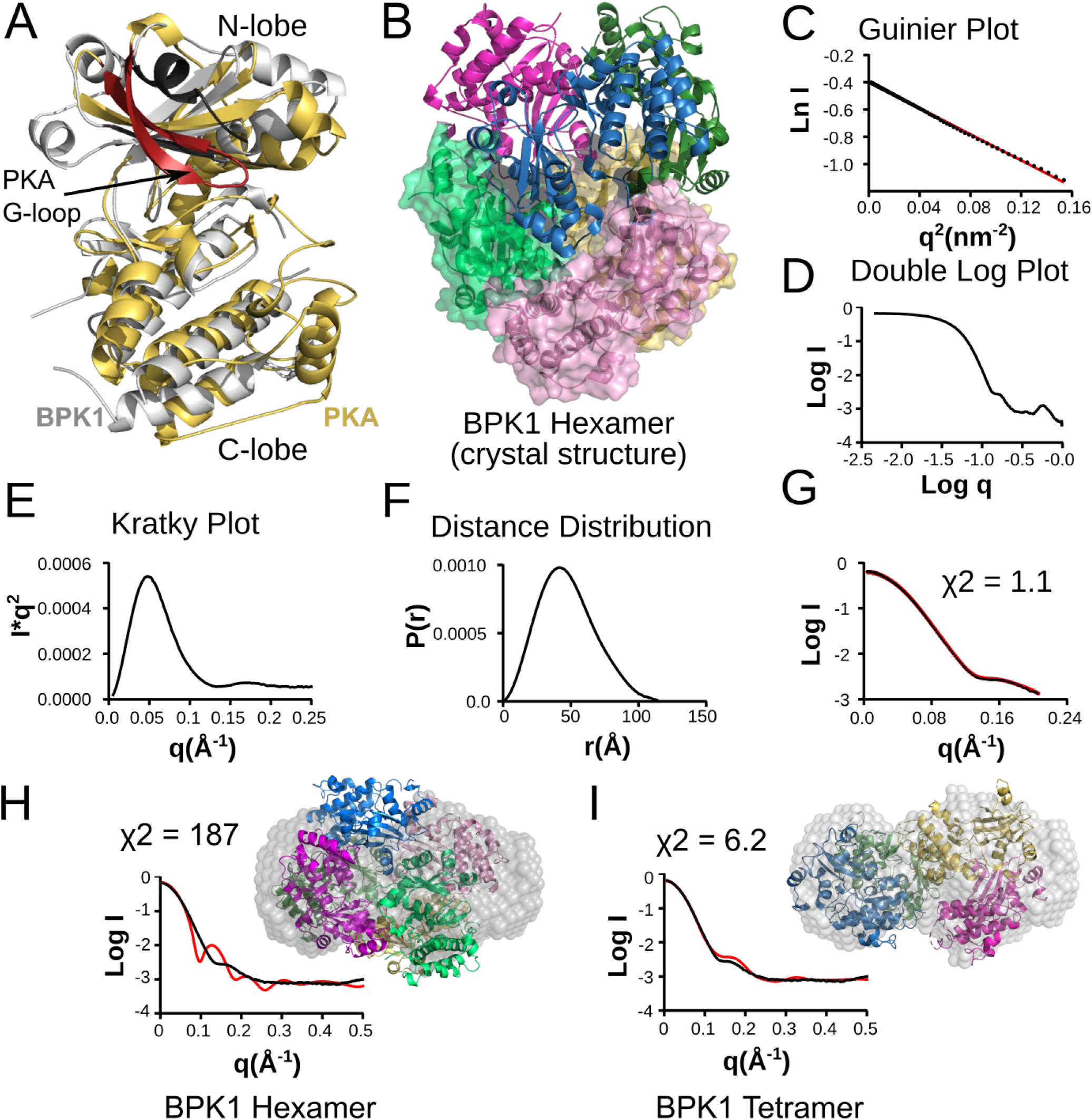
The WNG family pseudokinase TgBPK1 is a homo-oligomeric protein. (A) The structures of human PKA1 and TgBPK1 are represented in yellow and gray respectively. The Gly-loop of PKA1 is represented in orange and the corresponding region in TgBPK1 is colored slate (adapted from (Beraki et al., 2019)). (B) The asymmetric unit in the TgBPK1 crystal structure (PDB ID: 6M7Z (Beraki et al., 2019). Three subunits are shown as cartoon (blue, magenta and green) while other subunits are also shown as surfaces (yellow, pink, sea green). (C) Guinier plot [Ln (q) vs q2], (D) Double log plot [Log q vs. Log I(q)], (E) Kratky Plot [I*q^2^ vs q] (F) pairwise distance distribution P(r) plot for TgBPK1 in-solution SAXS data analysis. (G) The calculated model (black) from DAMMIN analysis was fitted with the experimental data (red) with a χ^2^ = 1.1. The molecular envelope generated for TgBPK1 by in-solution SAXS data analysis fitted with the TgBPK1 homohexamer (H) and homotetramer (I) with the corresponding graphs of their fit to experimental data. The tetramer showed better fit with χ^2^ value of 6.2, while the hexamer has χ^2^ of 187. The individual subunits in the tetramer are shown as blue, yellow magenta and green. The hexamer color scheme is same as the (B). The black line indicates the experimental scattering data and the red line indicates the pdb based scattering generated for fitting.

In the present work, we sought to determine the mechanism by which WNG1 activity is regulated. We found that another WNG-family member, BPK1, can exist in different multimeric states. WNG1, however, does not multimerize in solution and is therefore unlikely to be regulated in this manner. Instead, we found that WNG1 can autophosphorylate two residues in its C-lobe distinct from the activation loop, and that these sites are required for WNG1 activity. We also identified a series of basic residues that are also required for WNG1 activity and therefore appear to be recognizing the activating phosphosites. Structural modeling revealed a mechanism by which these regulatory salt bridges may stabilize the WNG1 C-lobe, analogous to activation loop recognition in canonical kinases.

## Results

### The WNG-family pseudokinase, BPK1, can adopt different multimeric states

One mechanism by which kinase activity can be regulated is through changes in the multimerization state. We had previously crystallized a WNG family member, the pseudokinase BPK1, and found that it forms homohexameric assembly in all crystal forms we obtained (Figure 1B). Notably, the BPK1 pseudoactive sites are oriented into the closed hexamer. Consistent with its multimerization, BPK1 has been proposed to play a structural role in *Toxoplasma* bradyzoite cyst wall formation (Buchholz et al., 2013). To confirm that the multimerization state is not an artifact of crystallization, we performed small-angle x-ray scattering (SAXS) on recombinantly purified BPK1. Initial analysis of the BPK1 scatter indicated high sample monodispersity of a well-folded protein with an approximate Rg of 40 Å (Figures 1C-F, Table 1). We used the DAMMIN (Svergun, 1999) program to obtain a molecular envelope for the most probable in-solution state of the BPK1. The resulting model showed good congruence with the experimental scattering values with a χ^2^ = 1.1 (Figure 1G). While the predicted molecular envelope was much larger than a BPK1 monomer, it was not large enough to contain the BPK1 hexamer from the crystal structure. Instead, the molecular weight calculated from the volume of correlation (Vc; Table 1) was consistent with a BPK1 tetramer. Additionally, while the crystallographic hexamer fit poorly in the calculated envelope (χ^2^ = 187; Figure 1H), the generated tetramer of BPK1 fit well in the molecular envelope (χ^2^ = 6.2; Figure 1I). Importantly, there is no tiling arrangement by which a tetrameric state can be arranged to form closed hexamers; to form the hexameric contacts we observed in the crystal, the BPK1 tetramer must first dissociate. Therefore, BPK1 can adopt different multimerization states.

**Table 1:** SAXS Parameters for BPK1.

| Method | Parameter |  |
| --- | --- | --- |
| Guinier Analysis | Rg | 38.49 |
| Indirect fourier transformation | Dmax | 114.55 |
| Distance Distribution | Rg | 41.9 |
| Porod Volume ($\text{\AA}^3$) | | 206804 |
| Molar Mass determination by Vc (kDa) | Mw | 148.24 |
| Theoretical Mw of Monomer (kDa) |  | 35.95 |

### WNG1 does not oligomerize in solution

Because the closed hexamer state of BPK1 would be unable to interact with substrate, we reasoned that such a multimeric state may be an inhibited state conserved among WNG family members. We therefore sought to determine if the WNG family member and an active kinase, WNG1, was also able to adopt similar multimerization states. We bacterially expressed the WNG1 kinase domain (residues 265-591) as a His_6_-SUMO fusion and purified the protein. We then compared the size exclusion chromatography (SEC) profile of WNG1 with that of a monomeric kinase, ERK7 (residues 2-350) that had been similarly tagged (Figure 2A). Both the WNG1 and ERK7 fusion proteins are of similar molar mass (51.4 kDa – WNG1 and 54.4 kDa – ERK7) and exhibit similar retention times on the SEC column. As the retention time for WNG1 was consistent with its monomeric molecular weight, WNG1, unlike BPK1, does not appear to form tightly associated multimers. However, we could not rule out that WNG1 formed transient multimers that may affect kinase activity. Such regulation of enzymatic activity by multimerization would be evident as a concentration-dependence on kinase specific activity (*i.e*., cooperativity). We therefore assessed WNG1 kinase activity at a range of WNG1 concentrations (0.25 – 3 μM) using myelin basic protein (MBP) as a substrate. WNG1 specific activity was independent of kinase concentration (Figure 2B), leading us to conclude that WNG1 activation is not dependent on multimerization.

**Figure 2:**
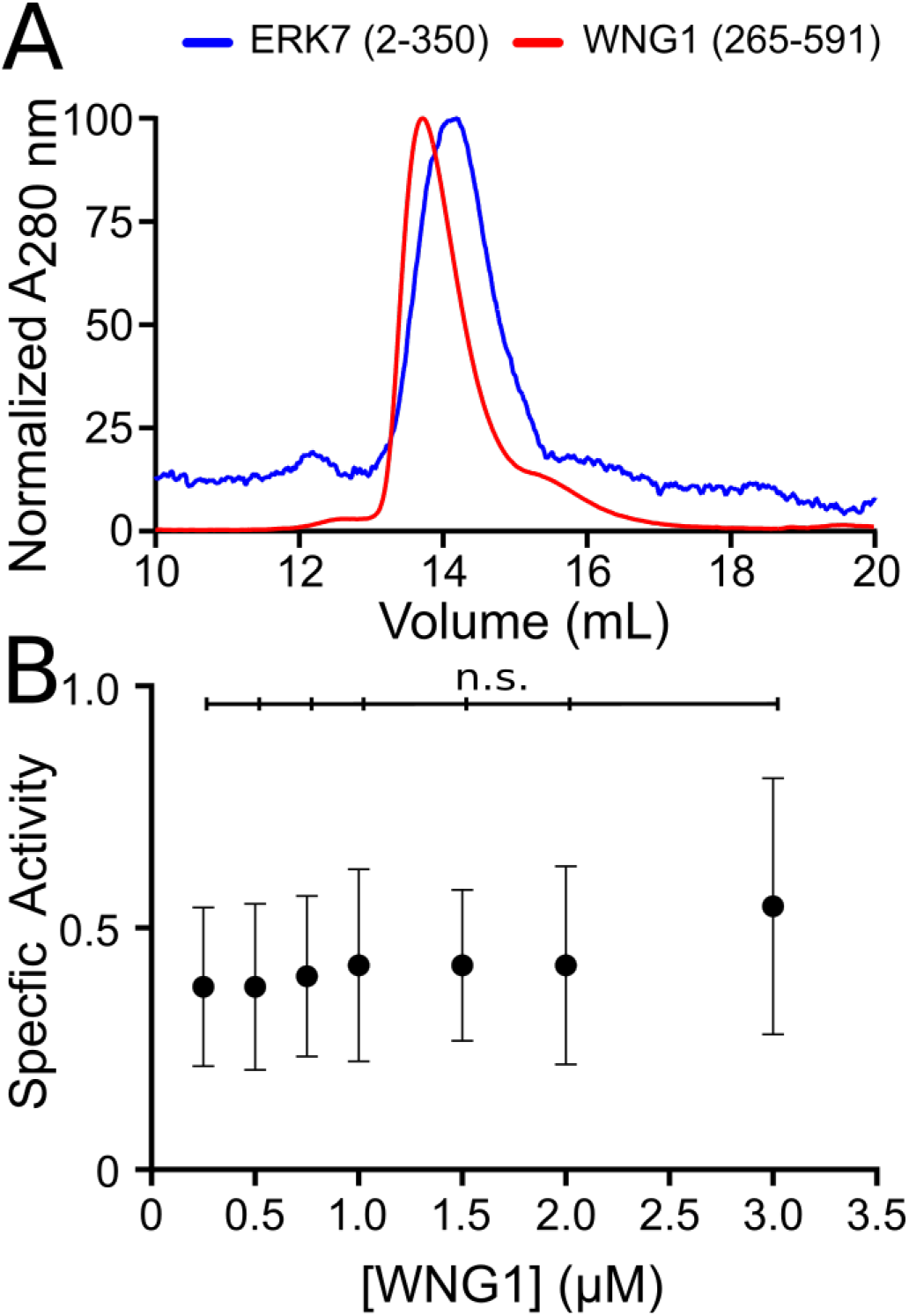
WNG1 is a monomer and its activity is concentration independent. (A) Comparison of size exclusion chromatography elution profiles for SUMO-ERK7(2-350) and SUMO-WNG1(265-591). (B) Kinase activity measurement for SUMO-WNG1(265-591) at the indicated enzyme concentrations. Significance was determined by One-way ANOVA followed by Tukey’s test; n.s. - not significant.

### WNG1 autophosphorylates residues not associated with regulation of other kinases

After ruling out multimerization of WNG1 as a likely regulatory mechanism, we tested whether WNG1 may be activated by phosphorylation. Recombinant WNG1 robustly autophosphorylates, suggesting it may autoactivate via this mechanism. We performed mass spectrometry to compare the phosphosites found on wild-type (WT) versus kinase-dead (D437S) WNG1. From these data, we identified residues Ser325, Ser349, Ser350, Ser480, Thr486 and Thr534 as phosphorylated. Since the residues identified in the case of WNG1 were present at unconventional positions (Figure 3A) we wanted to know if their phosphorylation was essential for its kinase activity. The first 3 sites are located in the N-lobe while the latter three are located in the C-lobe of WNG1 (Figure 3A,B). Strikingly, none of these residues are within the WNG1 activation loop, and to our knowledge, are not at sites that have been associated with activation of other kinases (Canagarajah et al., 1997; Xu et al., 2002; Zheng et al., 1993). To test whether phosphorylation of these phosphosites is essential for WNG1 activity, we mutated each of the identified phosphosites to Ala. Notably, WNG1 has one phosphorylatable residue, Thr466, in its activation loop (Figure 3A,B), though we did not identify it as phosphorylated in its activation loop. However, we had previously shown that WNG1 requires its HRD Arg436 for activity (Beraki et al., 2019), a site which normally recognizes phosphorylated Ser/Thr in a canonical kinase activation loop (Canagarajah et al., 1997; Zheng et al., 1993). To confirm Thr466 is not required for WNG1 activity, we also created a T466A mutant.

**Figure 3:**
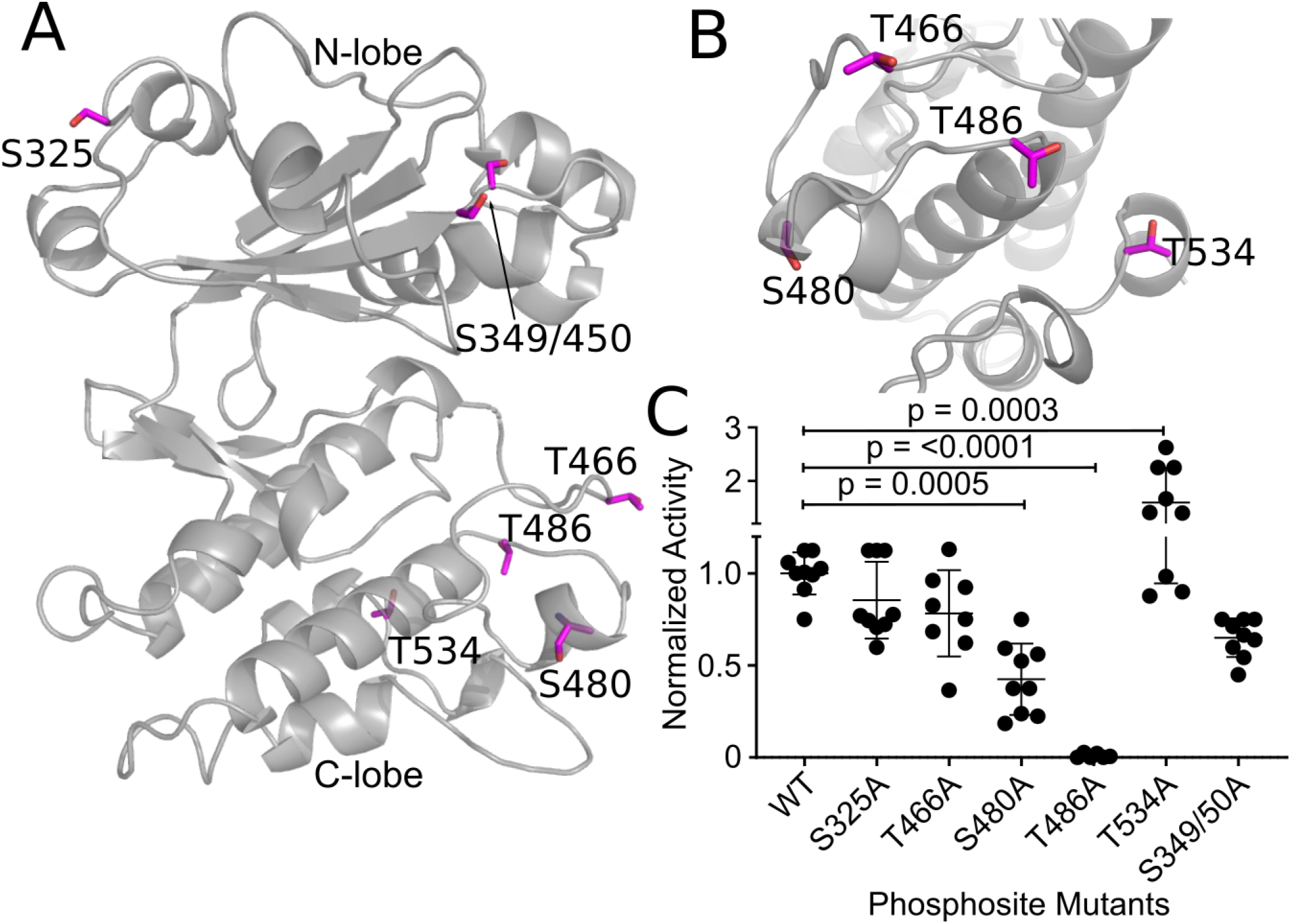
WNG1 is activated by phosphorylation. (A) WNG1 homology model generated from BPK1 structure is represented in gray color. The serine/threonine residues identified from the mass spectrometry are shown as sticks in magenta color. (B) Rotated (−90°) and zoomed view of the WNG1 C-lobe serine/threonine phosphosites (magenta sticks). (C) Kinase activity of WNG1 protein with non-phosphorylatable (alanine) mutants of the Ser/Thr phosphosites. Significance was determined by One-way ANOVA followed by Dunnett’s multiple comparisons test.

We evaluated the *in vitro* kinase activities of each mutant. Mutation of the three N-lobe sites did not significantly affect kinase activity (Figure 3C). Thus phosphorylation in the N-lobe of WNG1 appears dispensable for its kinase activity. We also observed no significant change in activity upon mutation of Thr466, confirming that WNG1 is not regulated by a canonical activation loop mechanism. Somewhat surprisingly, mutation of Thr534 increased the specific activity of the kinase (Figure 3C), suggesting that this residue is either a site of negative regulation or its mutation stabilizes the kinase. On the other hand, the Ser480 mutant had ~40% activity of wild-type WNG1 (Figure 3C). Most strikingly, mutation of Thr486 rendered WNG1 completely inactive. Taken together, these data suggest that WNG1 requires phosphorylation on non-canonical sites on its C-lobe for kinase activity.

### Basic residues in proximity of T486 are essential for WNG1 activity

As strongly electronegative residues, phosphosites are typically recognized by basic patches of Lys and Arg (Canagarajah et al., 1997; Russo et al., 1996; Zheng et al., 1993). We therefore sought to identify basic residues that may recognize the WNG1 activating phosphosites Ser480 and Thr486. We generated a homology model of the WNG1 structure using TgBPK1 as a template. We used this model to identify basic residues with side chains within 4 Å of Thr486 or Ser480 (Figure 4). Consistent with its published mutant effect on kinase activity, we found that the HRD Arg436 is within 4 Å of Thr486, and may therefore recognize its phospho-state. In addition to Arg436, four other Arg/Lys are within range of the phosphosites to potentially form salt bridges (Figure 4A). Notably, these basic residues are each conserved in WNG1 kinases from other species (Figure 5). To test if these basic residues were essential for kinase activity, we mutated each individually. We expressed and purified each mutant protein and tested its *in vitro* kinase activity. Mutants of Arg436 and Arg532 were completely inactive (Figure 4B), consistent with potential roles in coordinating a phosphorylated Thr486. Lys488 is also nearby Thr486, though its mutation to Ala resulted in only a minor 40% reduction in activity (Figure 4B). Lys522 and Arg523 are each nearby S480. While mutation of Lys522 did not affect WNG1 kinase activity, the Arg523 mutant showed a partial reduction in activity (Figure 4B). Taken together, these data indicate that Thr486 phosphorylation is essential to activity and is coordinated by Arg436 and Arg532, while Ser480 is a secondary regulatory site and may be coordinated by Arg523.

**Figure 4:**
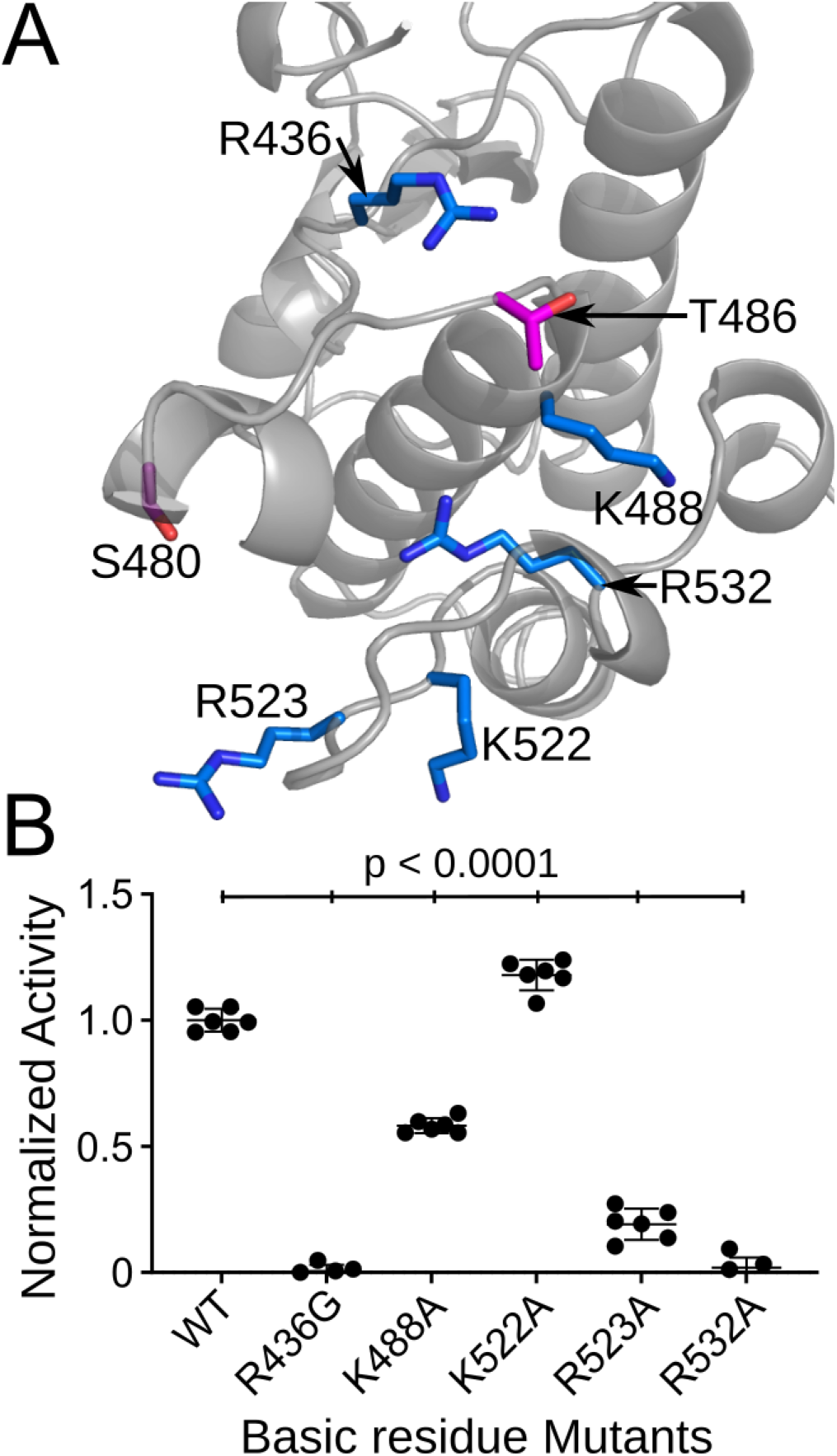
Basic residues in proximity of the phosphosites are important for WNG1 kinase activity. (A) C-lobe of the WNG1 homology model showing the basic amino acids (represented as magenta sticks) within 10 Å of S480 and T486 (represented as green sticks). (B) Kinase activity measurement of the WNG1 protein with alanine mutation at R436, K488, K522, R523 and R532. Significance was determined by One-way ANOVA followed by Dunnett’s multiple comparisons test.

**Figure 5:**
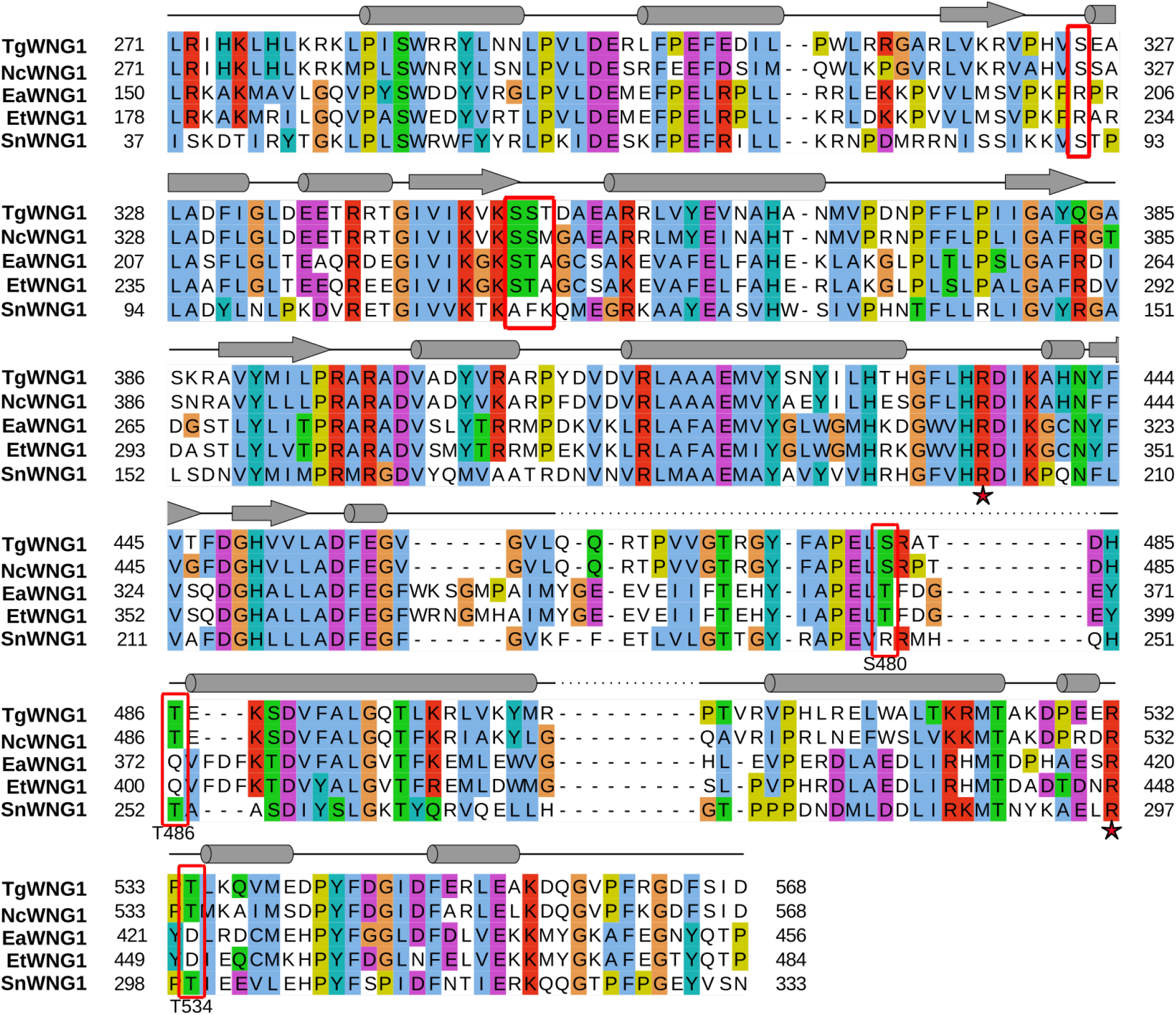
Sequence alignment of WNG1 showing phosphosites. Sequence alignment of the WNG1 kinase domain and its orthologs from selected coccidian parasites is shown with ClustalW color pattern. The residues that are autophososphorylated in the recombinant protein are highlighted in red rectangles. The Arg residues R436 and R532 that are near phosphosites and are essential for WNG1 activity are marked with red star. The predicted secondary structure is represented on top of the corresponding WNG1 sequence; helices as cylinders and β-sheet is represented by arrows (Tg – *Toxoplasma gondii*, Nc – *Neospora caninum*, Ea – *Eimeria acervulina*, Et – *Eimeria tenella*, Sn – *Sarcocystis neurona*).

To understand the salt bridge network formed by these interactions, we mapped the basic residues on the WNG1 model with phosphorylated Thr486 and Ser480 (Figure 6). In the model, the helix containing pThr486 was moved away from the putative activation loop to correctly orient pThr486, Arg436, and Arg532 for salt bridge formation (Figure 6A, 6B). The Thr486 in the unphosphorylated WNG1 is otherwise away from these basic residues (Figure 6C). Similarly, the pSer480 phosphoryl group can be coordinated by Arg523 (Figure 6D), while the Ser480 hydroxyl group will not be proximal for a salt bridge with Arg523 (Figure 6E). Upon activation of WNG1, the formation of these salt bridges may result in the movement of the helix containing pThr486 away from the putative activation loop (Figure 6A). The salt bridge interaction network that stabilizes the new position of pThr486 containing helix is localized in the C-lobe of the WNG1 (Figure 7A). Analogous, but distinct salt bridge interactions of the phosphorylated residues in canonical kinases stabilize the activation loop and parts of both the N- and C-lobes (*e.g.* PKA and ERK2; Figure 7). In contrast, our model suggests that for WNG1, the net effect of activation appears to be C-lobe movements that open up the activation site, providing WNG1 with a wider catalytic site.

**Figure 6:**
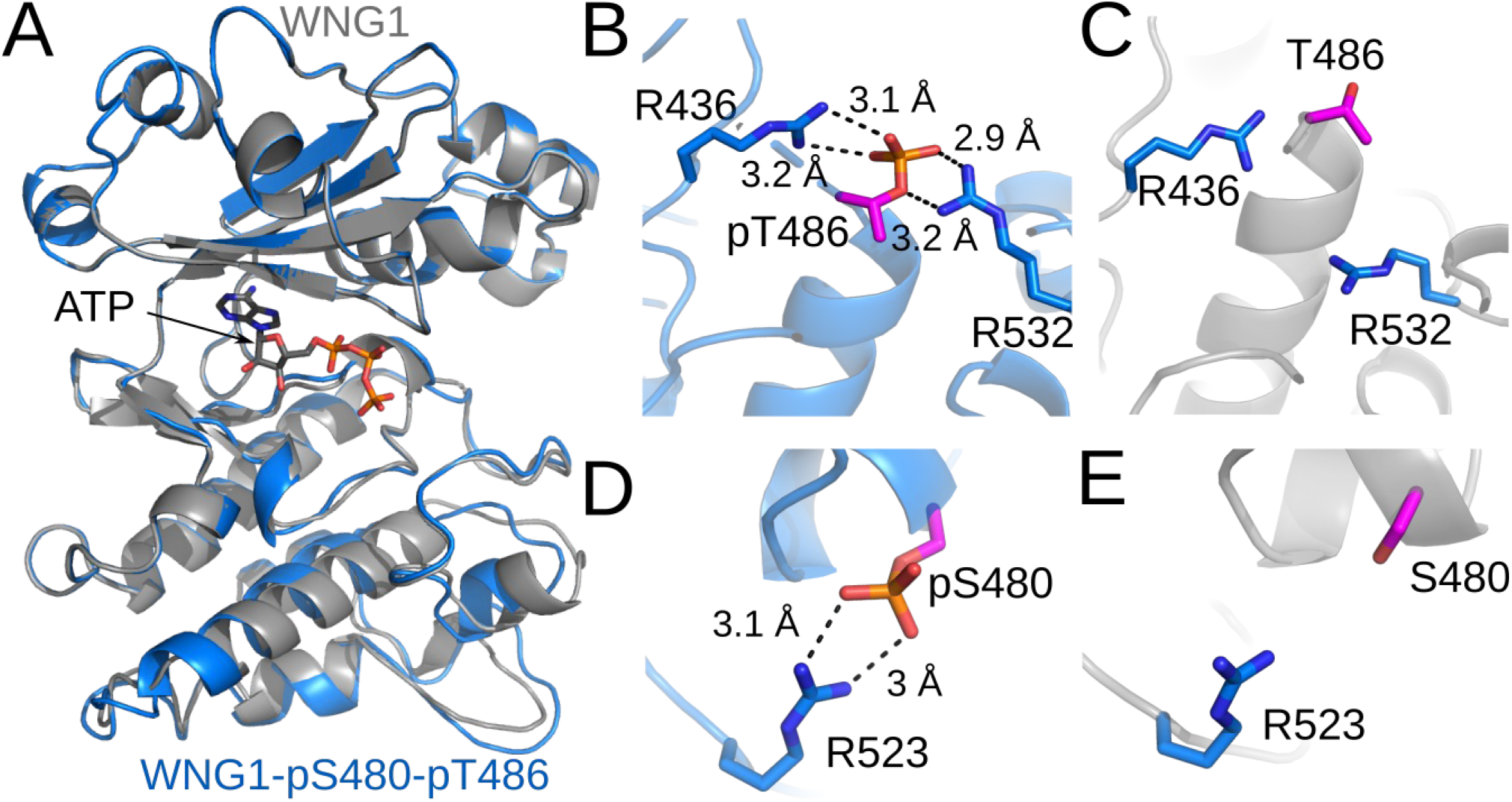
WNG1 activating phosphorylation stabilizes the C-lobe. (A) Superimposition of the WNG1 model on the WNG1-pS480-pT486 model represented in gray and blue color respectively. The ATP is shown gray color. Phosphosites, T486 and S480 are shown in magenta and corresponding basic residues (R436, R532, and R523) are shown in blue stick representation for pT486-pS480-WNG1 (B, D) and WNG1 (C, E). Salt bridges are indicated as back dotted lines.

**Figure 7:**
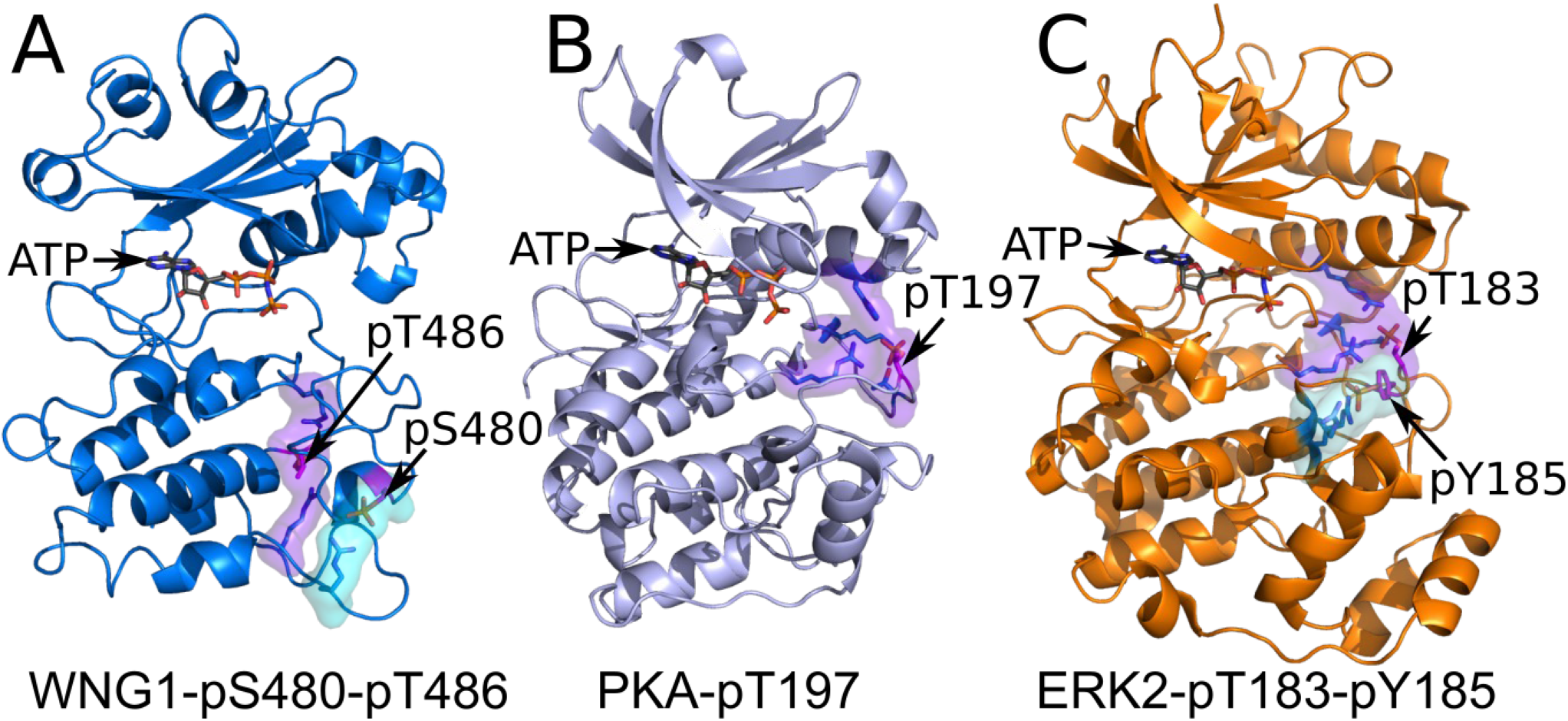
Comparison of salt bridge networks stabilizing activating phosphosites. The residues that form salt bridge interaction with the phosphorylated residue (magenta) are shown as blue sticks. The primary and secondary phosphosite interaction networks are represented in purple and cyan surface respectively. The ATP is in gray color. (A) WNG1-pS480-pT486 is represented in blue color. (B) PKA1-pT197 is represented in light blue color. For better visibility some portions of PKA have been omitted (C) ERK2-pT183-pY185 is represented in orange.

## Discussion

We have identified that the divergent kinase WNG1 is activated by autophosphorylation of residues Ser480 and Thr486. Upon phosphorylation, these two residues are coordinated by basic residues in their proximity. The formation of these salt bridges is essential and stabilizes the C-lobe in a new conformation (Figures 6-7). In contrast to the canonical kinases where the activation loop phosphorylation results in stabilized N and C lobes, WNG1 phosphorylation likely results in major conformational changes only within its C-lobe as we did not identify an activating phosphosite on the α-C helix.

The substrates of a canonical kinase are typically phosphorylated on flexible loops (Iakoucheva, 2004). The kinase activation loop forms β-strand-like interactions with the substrate backbone to position it correctly in the active site (Hubbard, 1997; Lowe, 1997). For this reason, recognition of activation loop phosphorylation is often critical to docking of substrate. Our data indicate that the WNG1 HRD Arg436 does not coordinate the activation loop to activate the kinase as occurs in canonical kinases. Consistent with this idea, WNG1, appears to phosphorylate the majority of its substrates on or proximal to α-helices (Beraki et al., 2019). As α-helices are characterized by a completed hydrogen-bond network along the sequence (Pauling et al., 1951), the backbone would not be available for interaction with the activation loop. Taken together, these data suggest that WNG1 must recognize its substrates in a manner distinct from a canonical kinase (Beraki et al., 2019), though further structural work is required to define that mechanism.

Several WNG1 substrates tubulate membranes upon integration (Mercier et al., 2002; Coppens et al., 2006; Lopez et al., 2015), which would be expected to be harmful to parasite if allowed to target incorrectly, such as to the parasite Golgi or plasma membranes. How WNG1 activity is controlled *in vivo* is therefore an important and outstanding question. We have previously demonstrated WNG1 is inactive while trafficking through the parasite secretory system (Beraki et al., 2019). It is likely that WNG1 is maintained inactive until it comes into proximity of the PV membrane to prevent premature insertion of substrates into the parasite plasma membrane. In addition, WNG1 is primarily associated with the PV membrane after secretion, though, unlike its substrates, the kinase is not an integral membrane protein (Beraki et al., 2019).

We propose a model for *in vivo* activation of WNG1 in which the major requirement in WNG1 activation is the availability of ATP (Figure 8). Recombinant WNG1 is active *in vitro* and robustly autophosphorylates, suggesting it does not require another kinase for its activation. Because the K_M,ATP_ for WNG1 (~500 μM) is relatively high compared to most typical kinases (Beraki et al., 2019), surrounding ATP levels must approach millimolar concentration for reasonable activity. ATP concentration in the PV lumen is thought to approximate the 2-5 mM concentration of the host cell due to diffusion through a pore in the PV membrane (Gold et al., 2015; Schwab et al., 1994). Regardless, ATP levels within the PV are clearly sufficient for WNG1 activity, as we have previously demonstrated (Beraki et al., 2019). However, *Toxoplasma* is auxotrophic for purines (Perrotto et al., 1971) and secretes highly active NTPases into the PV lumen to convert ATP to adenosine for parasite uptake at its plasma membrane (Sibley et al., 1994). Taken together, these data suggest a strong gradient of ATP concentration in the PV lumen where levels are lowest near the parasite plasma membrane and highest near pores in the PV membrane. We therefore propose that WNG1 activates as it approaches the PV membrane, after which it becomes tightly associated with an as-yet unknown integral membrane partner. The activated WNG1 then phosphorylates GRA proteins and induces their integration into the PV membrane.

**Figure 8:**
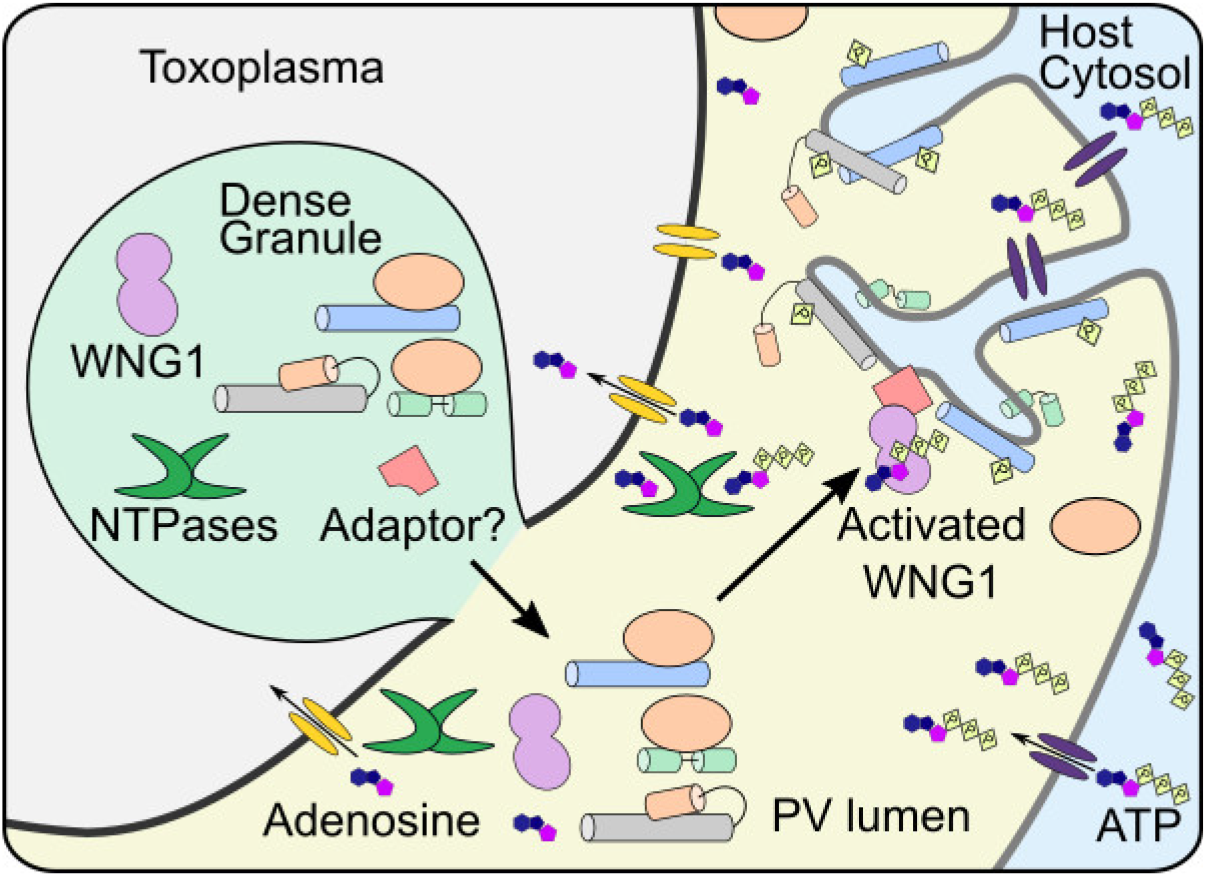
Model for activation of WNG1 *in vivo*. The dense granule contains WNG1, NTP hydrolases, other GRA proteins (blue, gray, green cylinders), their chaperone (orange) and the adaptor for WNG1 membrane integration. Upon release into the PV, the NTP hydrolases convert the ATP to adenosine for parasite uptake, reducing the ATP concentration in the lumen. The released WNG1 diffuses to the PV membrane where it gets autoactivated in the presence of high ATP levels. The activated WNG1 adheres to the PV membrane with the help of the adapter protein. Activated WNG1 phosphorylates GRAs resulting in their insertion into the PV membrane and formation of the IVN. The adenosine transport channel is represented in yellow while the ATP transporter in the PV membrane is represented in purple.

## Materials and Methods

### PCR and Mutagenesis

The WNG1(265-591) and BPK1(61-377) cDNA was previously cloned in the pET28a vector with the SUMO tag (Beraki et al., 2019). Mutagenesis was performed by Phusion QuikChange protocol (Xia et al., 2015) using Phusion polymerase (New England Biolabs). The primers are listed in table 2.

**Table 2:**
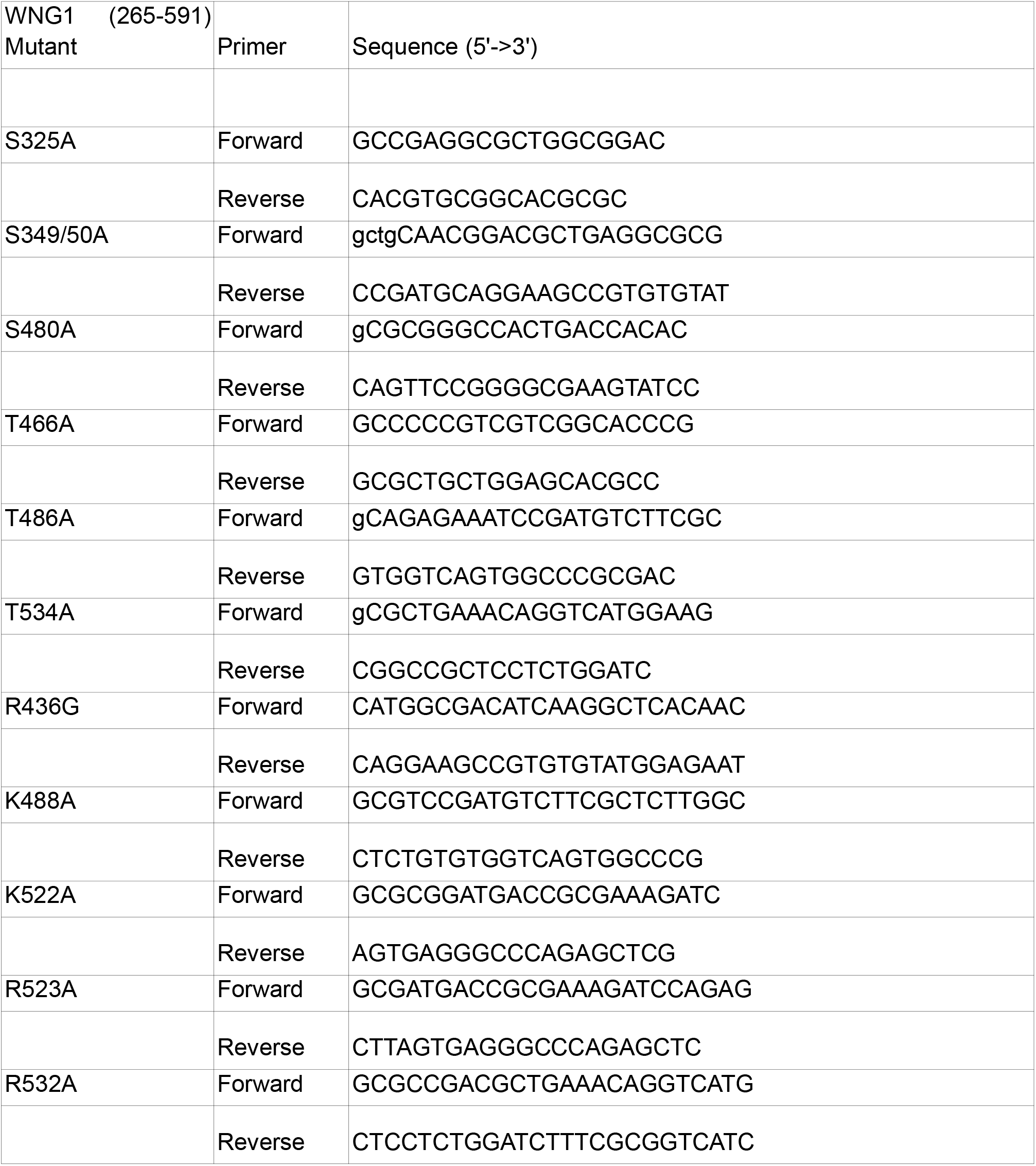
Primers used for WNG1 mutagenesis.

### Protein Purification

WNG1 wildtype, mutants, and BPK1 (61-377) were purified as per the protocol mentioned in (Beraki et al., 2019). All the proteins were expressed in Rosetta 2 (DE3) cells with a His_6_-SUMO tag overnight at 16 degrees post-induction with 500 mM IPTG. The WNG1 expressing cells were resuspended in 20 mM Tris HCl pH 7.5, 500 mM NaCl and 15 mM Imidazole (Resuspension buffer) and lysed by sonication. The lysate was centrifuged at 48k rcf and bound to Ni-NTA agarose beads (Qiagen). The beads were washed with the resuspension buffer and eluted with 20 mM Tris HCl pH 7.5, 500 mM NaCl and 300 mM Imidazole. The elution buffer was exchanged to 20 mM HEPES pH 7.5, 300 mM NaCl, 5 mM MgCl_2_ and 10% glycerol during concentration. The concentrated protein was further purified by size exclusion chromatography (SEC) first using Sephacryl 12 HR 16/60 and then Superdex 200 pg 16/60 columns. The final protein was concentrated and flash frozen for long-term storage in 20 mM HEPES pH 7.5, 300 mM NaCl, 5 mM MgCl_2_ and 10% glycerol buffer. The affinity chromatography for BPK (residues 61-377) protein was identical to the WNG1 protein preparation. The BPK1 protein was digested with ULP1 post Ni-NTA elution and anion exchange chromatography was performed to remove the SUMO Tag. SEC was performed as the final stage of purification using 20mM HEPES pH 7.5, 100mM NaCl as the buffer. The final protein was concentrated and flash frozen and stored until further use.

### Mass spectrometry for phosphosite identification

The recombinant wildtype WNG1 and the kinase dead D437S mutant recombinant protein was loaded on BioRad Mini-PROTEAN Precast gel. The gel was stained with Gel-code Blue (Thermo Fisher) and the WNG1 bands were cut out into small pieces and transferred to fresh tubes. The gel samples were digested overnight with trypsin (Pierce) following reduction and alkylation with DTT and iodoacetamide (Sigma–Aldrich). The samples then underwent solid-phase extraction cleanup with an Oasis HLB plate (Waters) and the resulting peptides were injected onto an Orbitrap Fusion Lumos mass spectrometer coupled to an Ultimate 3000 RSLC-Nano liquid chromatography system. Samples were injected onto a 75 um i.d., 50-cm long EasySpray column (Thermo) and eluted with a gradient from 0-28% buffer B over 60 min at 250 nL/min. Buffer A contained 2% (v/v) ACN and 0.1% formic acid in water, and buffer B contained 80% (v/v) ACN, 10% (v/v) trifluoroethanol, and 0.1% formic acid in water. The mass spectrometer operated in positive ion mode with a source voltage of 2.2 kV and an ion transfer tube temperature of 275 °C. MS scans were acquired at 120,000 resolution in the Orbitrap and up to 10 MS/MS spectra were obtained in the ion trap for each full spectrum acquired using higher-energy collisional dissociation (HCD) for ions with charges 2-7. Dynamic exclusion was set for 25 s after an ion was selected for fragmentation.

Raw MS data files were analyzed using Proteome Discoverer v2.2 (Thermo), with peptide identification performed using Sequest HT searching against the e. coli protein database from UniProt along with the wild type and kinase dead sequences of ROP35. Fragment and precursor tolerances of 10 ppm and 0.6 Da were specified, and three missed cleavages were allowed. Carbamidomethylation of Cys was set as a fixed modification, with oxidation of Met, methylation of Lys and Arg, demethylation of Lys and Arg, trimethylation of Lys, acetylation of Lys, and phosphorylation of Ser, Thr, and Tyr set as a variable modification. The false-discovery rate (FDR) cutoff was 1% for all peptides.

### In Vitro Kinase Assays

The kinase assays were performed using 2 μM of SUMO-WNG1 (residues 265-591) protein (mutant and wildtype), 20 mM HEPES pH 7.5, 300 mM NaCl, 5 mM MgCl2, 10% glycerol, 1mM DTT, 1mg/mL BSA, and 200 μM cold ATP. Hot ATP equivalent of 10 μCi and 5 mM MBP (substrate) was added at the end and incubated at 30°C for 2 hours. The reaction was stopped by adding 6x SDS loading dye. The kinase activity of the proteins was visualized or quantified by running the reactions on 15% SDS/PAGE. The gel was stained with coomassie stain and the MBP bands were cut out and the radioactivity was measured using a scintillation counter. The data were analyzed using Graphpad Prism 9.

### Small angle X-ray Scattering

Purified BPK1 protein was used for SAXS analysis at the Advanced Proton Source (APS), Lemont, USA. The protein was diluted from a stock concentration of 305 μM in 20mM HEPES pH 7.5, 100mM NaCl and the scattering data was collected at the ID12 beamline. The SAXS dataset was analyzed using the ATSAS software package (Manalastas-Cantos et al., 2021). DAMMIN (Svergun, 1999) was used for the construction of the Ab-initio molecular envelope. Five independent models were generated in the DAMMIN run and the theoretical I(q) was compared with the experimental I(q) for each of them. The model with the lowest Chi-square (χ^2^) value was used for fitting the BPK1 tetramer model using SUPALM (Konarev et al., 2016). SASREF (Petoukhov and Svergun, 2005) from the ATSAS suite was used for making a P2 symmetry-based BPK1 tetramer. We used a monomer from the BPK1 crystal structure (PDB ID - 6M7Z) (Beraki et al., 2019) for the tetramer reconstruction. SAXS data were deposited in SASBDB (accession: SASDNT5).

### Structure Modeling

The WNG1 homology model was generated using BPK1 crystal structure (PDB:6M7Z) as template in Modeller v9.14 (Beraki et al., 2019; Sali and Blundell, 1993). The pS480-pT486-WNG1 model was generated in COOT from the WNG1 model. Real-time refinement was used to optimize the geometry of the model in COOT.

### Figure generation

All figures were created in Inkscape. PyMOL (Schrödinger, LLC, 2015) was used for the analysis of structural models and the generation of ray-traced figures. Graphpad Prism was used to plot kinase activity graphs and to perform statistical analyses.

## Acknowledgements

We thank the Structural Biology Lab at UT Southwestern for their help with SAXS data collection. The SAXS measurements were performed at beamline 12-ID-B of the Advanced Photon Source, a U.S. Department of Energy (DOE) Office of Science User Facility, operated for the DOE Office of Science by Argonne National Laboratory under Contract No. DE-AC02-06CH11357, and was supported by the US DOE Office of Science, Office of Basic Energy Sciences. We acknowledge the University of Texas Southwestern medical center proteomics core facility for mass spectrometry analysis. We thank William O’Shaugnessy for helpful comments on the manuscript. M.L.R. acknowledges funding from the Welch Foundation (I-2075-20210327), National Science Foundation (MCB1553334), and NIH (AI150715).

## Data Availability Statement

SAXS data were deposited in SASBDB (accession: SASDNT5).

## Notes

### Competing Interest Statement

The authors have declared no competing interest.

## References

Abe, M.K., Kahle, K.T., Saelzler, M.P., Orth, K., Dixon, J.E., Rosner, M.R., 2001. ERK7 is an autoactivated member of the MAPK family. J. Biol. Chem. 276, 21272–21279. https://doi.org/10.1074/jbc.M100026200

Beraki, T., Hu, X., Broncel, M., Young, J.C., O’Shaughnessy, W.J., Borek, D., Treeck, M., Reese, M.L., 2019. Divergent kinase regulates membrane ultrastructure of the Toxoplasma parasitophorous vacuole. Proc. Natl. Acad. Sci. U. S. A. 116, 6361–6370. https://doi.org/10.1073/pnas.1816161116

Boulton, T.G., Nye, S.H., Robbins, D.J., Ip, N.Y., Radziejewska, E., Morgenbesser, S.D., DePinho, R.A., Panayotatos, N., Cobb, M.H., Yancopoulos, G.D., 1991. ERKs: a family of protein-serine/threonine kinases that are activated and tyrosine phosphorylated in response to insulin and NGF. Cell 65, 663–675.

Boulton, T.G., Yancopoulos, G.D., Gregory, J.S., Slaughter, C., Moomaw, C., Hsu, J., Cobb, M.H., 1990. An insulin-stimulated protein kinase similar to yeast kinases involved in cell cycle control. Science 249, 64–67.

Bradley, P.J., Ward, C., Cheng, S.J., Alexander, D.L., Coller, S., Coombs, G.H., Dunn, J.D., Ferguson, D.J., Sanderson, S.J., Wastling, J.M., Boothroyd, J.C., 2005. Proteomic Analysis of Rhoptry Organelles Reveals Many Novel Constituents for Host-Parasite Interactions in Toxoplasma gondii. J. Biol. Chem. 280, 34245–34258. https://doi.org/10.1074/jbc.M504158200

Buchholz, K.R., Bowyer, P.W., Boothroyd, J.C., 2013. Bradyzoite pseudokinase 1 is crucial for efficient oral infectivity of the Toxoplasma gondii tissue cyst. Eukaryot. Cell 12, 399–410. https://doi.org/10.1128/EC.00343-12

Canagarajah, B.J., Khokhlatchev, A., Cobb, M.H., Goldsmith, E.J., 1997. Activation Mechanism of the MAP Kinase ERK2 by Dual Phosphorylation. Cell 90, 859–869. https://doi.org/10.1016/S0092-8674(00)80351-7

Chao, L.H., Pellicena, P., Deindl, S., Barclay, L.A., Schulman, H., Kuriyan, J., 2010. Intersubunit capture of regulatory segments is a component of cooperative CaMKII activation. Nat. Struct. Mol. Biol. 17, 264–272. https://doi.org/10.1038/nsmb.1751

Coppens, I., Dunn, J.D., Romano, J.D., Pypaert, M., Zhang, H., Boothroyd, J.C., Joiner, K.A., 2006. Toxoplasma gondii sequesters lysosomes from mammalian hosts in the vacuolar space. Cell 125, 261–274. https://doi.org/10.1016/j.cell.2006.01.056

Davies, H., Bignell, G.R., Cox, C., Stephens, P., Edkins, S., Clegg, S., Teague, J., Woffendin, H., Garnett, M.J., Bottomley, W., Davis, N., Dicks, E., Ewing, R., Floyd, Y., Gray, K., Hall, S., Hawes, R., Hughes, J., Kosmidou, V., Menzies, A., Mould, C., Parker, A., Stevens, C., Watt, S., Hooper, S., Wilson, R., Jayatilake, H., Gusterson, B.A., Cooper, C., Shipley, J., Hargrave, D., Pritchard-Jones, K., Maitland, N., Chenevix-Trench, G., Riggins, G.J., Bigner, D.D., Palmieri, G., Cossu, A., Flanagan, A., Nicholson, A., Ho, J.W.C., Leung, S.Y., Yuen, S.T., Weber, B.L., Seigler, H.F., Darrow, T.L., Paterson, H., Marais, R., Marshall, C.J., Wooster, R., Stratton, M.R., Futreal, P.A., 2002. Mutations of the BRAF gene in human cancer. Nature 417, 949–954. https://doi.org/10.1038/nature00766

Eswaran, J., Patnaik, D., Filippakopoulos, P., Wang, F., Stein, R.L., Murray, J.W., Higgins, J.M.G., Knapp, S., 2009. Structure and functional characterization of the atypical human kinase haspin. Proc. Natl. Acad. Sci. 106, 20198–20203. https://doi.org/10.1073/pnas.0901989106

Gendrin, C., Mercier, C., Braun, L., Musset, K., Dubremetz, J.-F., Cesbron-Delauw, M.-F., 2008. Toxoplasma gondii uses unusual sorting mechanisms to deliver transmembrane proteins into the host-cell vacuole. Traffic Cph. Den. 9, 1665–1680. https://doi.org/10.1111/j.1600-0854.2008.00793.x

Gold, D.A., Kaplan, A.D., Lis, A., Bett, G.C.L., Rosowski, E.E., Cirelli, K.M., Bougdour, A., Sidik, S.M., Beck, J.R., Lourido, S., Egea, P.F., Bradley, P.J., Hakimi, M.-A., Rasmusson, R.L., Saeij, J.P.J., 2015. The Toxoplasma Dense Granule Proteins GRA17 and GRA23 Mediate the Movement of Small Molecules between the Host and the Parasitophorous Vacuole. Cell Host Microbe 17, 642–652. https://doi.org/10.1016/j.chom.2015.04.003

Hubbard, S.R., 1997. Crystal structure of the activated insulin receptor tyrosine kinase in complex with peptide substrate and ATP analog. EMBO J. 16, 5572–5581. https://doi.org/10.1093/emboj/16.18.5572

Hunter, T., 1995. Protein kinases and phosphatases: The Yin and Yang of protein phosphorylation and signaling. Cell 80, 225–236. https://doi.org/10.1016/0092-8674(95)90405-0

Iakoucheva, L.M., 2004. The importance of intrinsic disorder for protein phosphorylation. Nucleic Acids Res. 32, 1037–1049. https://doi.org/10.1093/nar/gkh253

James, C., Ugo, V., Le Couédic, J.-P., Staerk, J., Delhommeau, F., Lacout, C., Garçon, L., Raslova, H., Berger, R., Bennaceur-Griscelli, A., Villeval, J.L., Constantinescu, S.N., Casadevall, N., Vainchenker, W., 2005. A unique clonal JAK2 mutation leading to constitutive signalling causes polycythaemia vera. Nature 434, 1144–1148. https://doi.org/10.1038/nature03546

Knighton, D.R., Zheng, J., Eyck, L.F.T., Ashford, V.A., Xuong, N.-H., Taylor, S.S., Sowadski, J.M., 1991. Crystal Structure of the Catalytic Subunit of Cyclic Adenosine Monophosphate-Dependent Protein Kinase. Science. https://doi.org/10.1126/science.1862342

Konarev, P.V., Petoukhov, M.V., Svergun, D.I., 2016. Rapid automated superposition of shapes and macromolecular models using spherical harmonics. J. Appl. Crystallogr. 49, 953–960. https://doi.org/10.1107/S1600576716005793

Lacronique, V., Boureux, A., Della Valle, V., Poirel, H., Quang, C.T., Mauchauffé, M., Berthou, C., Lessard, M., Berger, R., Ghysdael, J., Bernard, O.A., 1997. A TEL-JAK2 Fusion Protein with Constitutive Kinase Activity in Human Leukemia. Science 278, 1309–1312. https://doi.org/10.1126/science.278.5341.1309

Lopez, J., Bittame, A., Massera, C., Vasseur, V., Effantin, G., Valat, A., Buaillon, C., Allart, S., Fox, B.A., Rommereim, L.M., Bzik, D.J., Schoehn, G., Weissenhorn, W., Dubremetz, J.-F., Gagnon, J., Mercier, C., Cesbron-Delauw, M.-F., Blanchard, N., 2015. Intravacuolar Membranes Regulate CD8 T Cell Recognition of Membrane-Bound Toxoplasma gondii Protective Antigen. Cell Rep. 13, 2273–2286. https://doi.org/10.1016/j.celrep.2015.11.001

Lowe, E.D., 1997. The crystal structure of a phosphorylase kinase peptide substrate complex: kinase substrate recognition. EMBO J. 16, 6646–6658. https://doi.org/10.1093/emboj/16.22.6646

Manalastas-Cantos, K., Konarev, P.V., Hajizadeh, N.R., Kikhney, A.G., Petoukhov, M.V., Molodenskiy, D.S., Panjkovich, A., Mertens, H.D.T., Gruzinov, A., Borges, C., Jeffries, C.M., Svergun, D.I., Franke, D., 2021. *ATSAS 3.0*: expanded functionality and new tools for small-angle scattering data analysis. J. Appl. Crystallogr. 54, 343–355. https://doi.org/10.1107/S1600576720013412

Manning, G., Whyte, D.B., Martinez, R., Hunter, T., Sudarsanam, S., 2002. The Protein Kinase Complement of the Human Genome. Science 298, 1912–1934. https://doi.org/10.1126/science.1075762

Mercier, C., Dubremetz, J.-F., Rauscher, B., Lecordier, L., Sibley, L.D., Cesbron-Delauw, M.-F., 2002. Biogenesis of nanotubular network in Toxoplasma parasitophorous vacuole induced by parasite proteins. Mol. Biol. Cell 13, 2397–2409. https://doi.org/10.1091/mbc.E02-01-0021

Moelling, K., Heimann, B., Beimling, P., Rapp, U.R., Sander, T., 1984. Serine-and threonine-specific protein kinase activities of purified gag-mil and gag-raf proteins. Nature 312, 558–561. https://doi.org/10.1038/312558a0

Oliver, A.W., Paul, A., Boxall, K.J., Barrie, S.E., Aherne, G.W., Garrett, M.D., Mittnacht, S., Pearl, L.H., 2006. Trans-activation of the DNA-damage signalling protein kinase Chk2 by T-loop exchange. EMBO J. 25, 3179–3190. https://doi.org/10.1038/sj.emboj.7601209

Pauling, L., Corey, R.B., Branson, H.R., 1951. The structure of proteins: Two hydrogen-bonded helical configurations of the polypeptide chain. Proc. Natl. Acad. Sci. 37, 205–211. https://doi.org/10.1073/pnas.37.4.205

Peixoto, L., Chen, F., Harb, O.S., Davis, P.H., Beiting, D.P., Brownback, C.S., Ouloguem, D., Roos, D.S., 2010. Integrative genomic approaches highlight a family of parasite-specific kinases that regulate host responses. Cell Host Microbe 8, 208–218. https://doi.org/10.1016/j.chom.2010.07.004

Perrotto, J., Keister, D.B., Gelderman, A.H., 1971. Incorporation of Precursors into *Toxoplasma* DNA. J. Protozool. 18, 470–473. https://doi.org/10.1111/j.1550-7408.1971.tb03356.x

Petoukhov, M.V., Svergun, D.I., 2005. Global Rigid Body Modeling of Macromolecular Complexes against Small-Angle Scattering Data. Biophys. J. 89, 1237–1250. https://doi.org/10.1529/biophysj.105.064154

Piala, A.T., Moon, T.M., Akella, R., He, H., Cobb, M.H., Goldsmith, E.J., 2014. Chloride sensing by WNK1 involves inhibition of autophosphorylation. Sci. Signal. 7, ra41. https://doi.org/10.1126/scisignal.2005050

Pike, A.C.W., Rellos, P., Niesen, F.H., Turnbull, A., Oliver, A.W., Parker, S.A., Turk, B.E., Pearl, L.H., Knapp, S., 2008. Activation segment dimerization: a mechanism for kinase autophosphorylation of non-consensus sites. EMBO J. 27, 704–714. https://doi.org/10.1038/emboj.2008.8

Rapp, U.R., Goldsborough, M.D., Mark, G.E., Bonner, T.I., Groffen, J., Reynolds, F.H., Stephenson, J.R., 1983. Structure and biological activity of v-raf, a unique oncogene transduced by a retrovirus. Proc. Natl. Acad. Sci. 80, 4218–4222. https://doi.org/10.1073/pnas.80.14.4218

Ray, L.B., Sturgill, T.W., 1988. Insulin-stimulated microtubule-associated protein kinase is phosphorylated on tyrosine and threonine in vivo. Proc. Natl. Acad. Sci. 85, 3753–3757. https://doi.org/10.1073/pnas.85.11.3753

Reese, M.L., Boyle, J.P., 2012. Virulence without catalysis: how can a pseudokinase affect host cell signaling? Trends Parasitol. 28, 53–57. https://doi.org/10.1016/j.pt.2011.12.004

Robbins, D.J., Zhen, E., Owaki, H., Vanderbilt, C.A., Ebert, D., Geppert, T.D., Cobb, M.H., 1993. Regulation and properties of extracellular signal-regulated protein kinases 1 and 2 in vitro. J. Biol. Chem. 268, 5097–106.

Russo, A.A., Jeffrey, P.D., Patten, A.K., Massagué, J., Pavletich, N.P., 1996. Crystal structure of the p27Kip1 cyclin-dependent-kinase inhibitor bound to the cyclin A-Cdk2 complex. Nature 382, 325–331. https://doi.org/10.1038/382325a0

Sali, A., Blundell, T.L., 1993. Comparative protein modelling by satisfaction of spatial restraints. J. Mol. Biol. 234, 779–815. https://doi.org/10.1006/jmbi.1993.1626

Schrödinger, LLC, 2015. The PyMOL Molecular Graphics System, Version 1.7.

Schwab, J.C., Beckers, C.J., Joiner, K.A., 1994. The parasitophorous vacuole membrane surrounding intracellular Toxoplasma gondii functions as a molecular sieve. Proc. Natl. Acad. Sci. U. S. A. 91, 509–513.

Sibley, L.D., Niesman, I.R., Asai, T., Takeuchi, T., 1994. Toxoplasma gondii: Secretion of a Potent Nucleoside Triphosphate Hydrolase into the Parasitophorous Vacuole. Exp. Parasitol. 79, 301–311. https://doi.org/10.1006/expr.1994.1093

Svergun, D.I., 1999. Restoring Low Resolution Structure of Biological Macromolecules from Solution Scattering Using Simulated Annealing. Biophys. J. 76, 2879–2886. https://doi.org/10.1016/S0006-3495(99)77443-6

Talevich, E., Mirza, A., Kannan, N., 2011. Structural and evolutionary divergence of eukaryotic protein kinases in Apicomplexa. BMC Evol. Biol. 11, 321. https://doi.org/10.1186/1471-2148-11-321

Tao, M., Salas, M.L., Lipmann, F., 1970. Mechanism of activation by adenosine 3’:5’-cyclic monophosphate of a protein phosphokinase from rabbit reticulocytes. Proc. Natl. Acad. Sci. U. S. A. 67, 408–414. https://doi.org/10.1073/pnas.67.1.408

Walker, J.E., Saraste, M., Runswick, M.J., Gay, N.J., 1982. Distantly related sequences in the alpha-and beta-subunits of ATP synthase, myosin, kinases and other ATP-requiring enzymes and a common nucleotide binding fold. EMBO J. 1, 945–951.

Xia, Y., Chu, W., Qi, Q., Xun, L., 2015. New insights into the QuikChangeTM process guide the use of Phusion DNA polymerase for site-directed mutagenesis. Nucleic Acids Res. 43, e12–e12. https://doi.org/10.1093/nar/gku1189

Xu, B., Min, X., Stippec, S., Lee, B.-H., Goldsmith, E.J., Cobb, M.H., 2002. Regulation of WNK1 by an Autoinhibitory Domain and Autophosphorylation. J. Biol. Chem. 277, 48456–48462. https://doi.org/10.1074/jbc.M207917200

Yang, Y., Ye, Q., Jia, Z., Côté, G.P., 2015. Characterization of the Catalytic and Nucleotide Binding Properties of the α-Kinase Domain of Dictyostelium Myosin-II Heavy Chain Kinase A. J. Biol. Chem. 290, 23935–23946. https://doi.org/10.1074/jbc.M115.672410

Zheng, J., Trafny, E.A., Knighton, D.R., Xuong, N., Taylor, S.S., Ten Eyck, L.F., Sowadski, J.M., 1993. 2.2 Å refined crystal structure of the catalytic subunit of cAMP-dependent protein kinase complexed with MnATP and a peptide inhibitor. Acta Crystallogr. D Biol. Crystallogr. 49, 362–365. https://doi.org/10.1107/S0907444993000423

